# Extracellular Vesicle exchange is favored by cell proximity

**DOI:** 10.1101/2022.05.10.491399

**Authors:** Federico Colombo, Erienne Grace Norton, Emanuele Cocucci

## Abstract

Extracellular vesicles (EVs) are biological nanovectors that retain information of the cell of origin and convey signals to recipient cells. Therefore, EVs are ideal platforms for the development of diagnostic tools and of bio-inspired drug delivery technologies. However, the dynamics of EV distribution in physiological conditions are still underexplored. Using an elegant series of experiments, including quantitative assays to define EV transfer and five dimension live cell imaging, we observe the release and internalization of EVs in real time and we demonstrate that EVs are mainly exchanged at the cell-cell interface. These observations prompt paradigm shifting consequences: first, EVs are mostly short-range intercellular vectors that influence adjacent cells; second, our data explain why increases in the internal pressure and permeability of the parenchyma, two hallmarks of inflammation and cancer, can facilitate EV escape from the damaged tissue. In conclusion, we provide experimental evidence supporting why EVs have great potential for the implementation of specific and sensitive liquid biopsy tests.

---

Extracellular vesicles (EVs) are membrane-delimited structures released by any cell that can navigate bodily fluids^1^. EVs contain molecules such as proteins and nucleic acids that have stimulatory potential and can influence multiple cellular functions, including cell motility and development, as well as contribute to disease progression^2–4^. Since the content of EVs is representative of their cell of origin, EVs released by diseased tissues carry cargoes that mirror the overall complexity of the underlying pathological process, representing a potential source of information for diagnostic and prognostic clinical use in cancer^5–7^, neurodegenerative diseases^8–10^, or other conditions for which biopsies are the standard of care but frequent repetition of the procedure is impractical due to invasiveness and cost^11^. However, the dynamics of EV distribution are poorly understood. For example, experimental and clinical cancer models show an increase of circulating EVs, but the biological basis for this observation is not yet explained^12^. Therefore shedding light on the biophysical rules that define the dynamics of EV exchange can provide essential information to exploit EVs as biomarkers and nanovectors for drug delivery. Since tissues are formed by densely packed arrays of cells, we hypothesized that cell density has a key role in controlling EV distribution by influencing EV release and/or diffusion.

To investigate the role of cell density in EV release, we plated SUM159-PT (SUM159) cells, a model of metastatic breast cancer^13^, at increasing concentrations. To avoid the confounding contribution of EVs derived from fetal bovine serum (FBS), we cultured the cells in Opti-MEM, which sustains cell growth in the absence of serum. After 24 hours, we collected the supernatants and processed them for Nanoparticle Tracking Analysis (NTA) to quantify EV concentration^14^. While the absolute number of EVs retrieved in the supernatant increased with increasing cell density, the number of EVs released per cell decreased significantly (Fig 1a). Increasing cell density did not significantly modify the size of the EVs, which maintained a mode diameter distribution between 129 and 152 nm (Figure 1b,c). We also observed the typical cup shape of EVs^15^ when the samples were subjected to transmission electron microscopy analysis (Figure 1d,e). Slot-blot analysis of the recovered media showed that the signals of the tetraspanins CD9 and CD63, well-known EV markers, followed a similar trend (Figure S1a-c). Since CD9 is a marker of ectosomes and CD63 of exosomes^16,17^, this result suggests that either EVs budding from the plasma membrane or EVs released upon multivesicular body fusion are influenced by cell density. A possible explanation is that sparse cells expose more surface to the extracellular space for EV budding than densely packed cells (Figure 1f).

**Figure 1.**
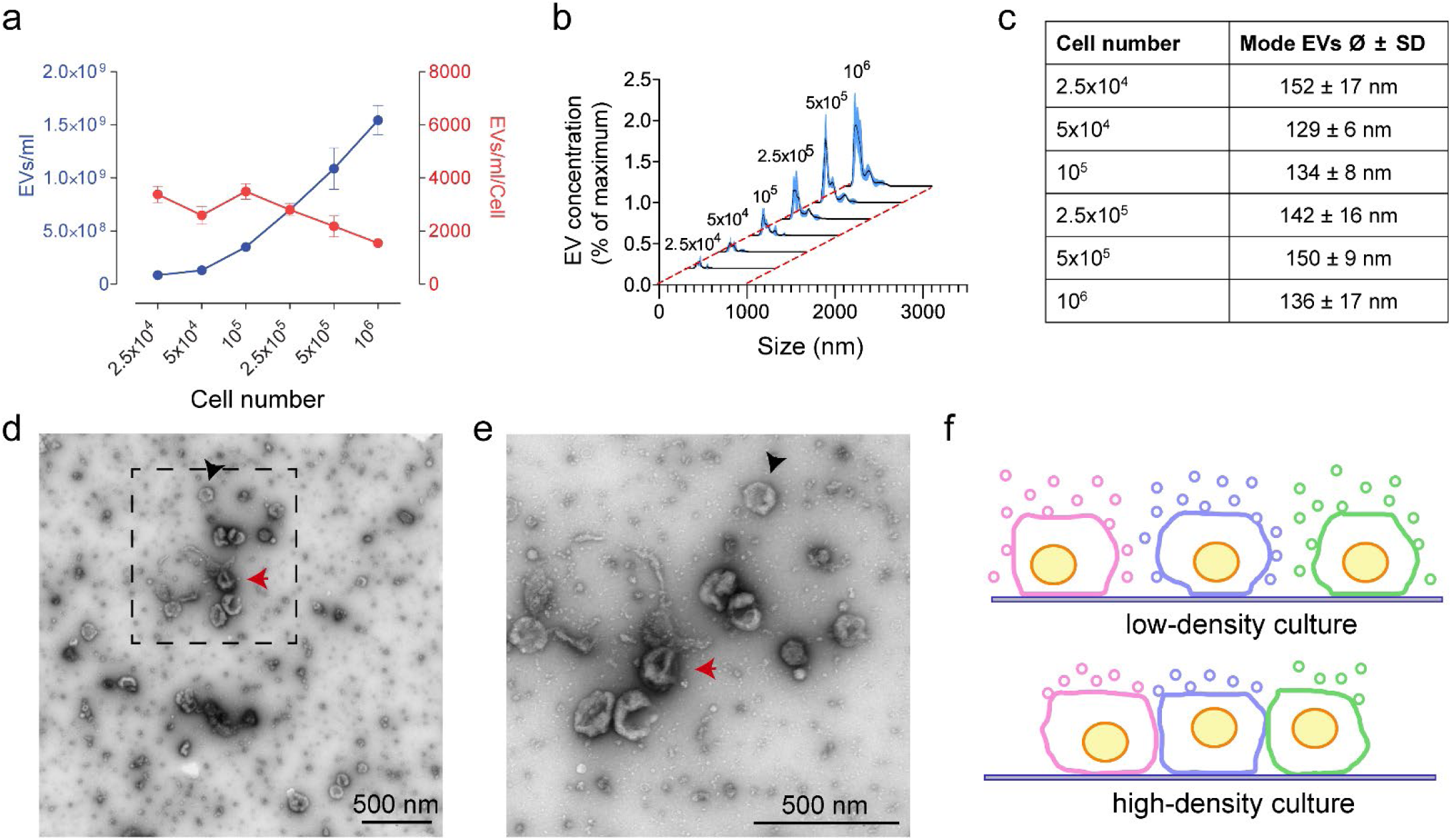
The number of EVs collected in the supernatant is inversely proportional to cell density. **a-c**, SUM159 cells were plated (6-well plate; well surface = 9.6 cm^2^) at increasing densities in Opti-MEM for 24 hours. After removing cell debris from the supernatants of each sample by centrifugation (5,000 x g for 10 minutes), both the EV concentration and the size distribution were assessed by Nanoparticle Tracking Analysis (NTA). **a**, While EV concentration increases with the number of cells (blue line), the number of EVs released per cell is inversely proportional to cell density (red line). The graph reports the mean ± SEM of three independent experiments. **b**, Size frequency distributions (mean = black line; SD = blue area) of EVs collected from cells plated at increasing numbers (indicated on top of each peak). Increasing cell density did not significantly influence the distribution of EV sizes. As tabulated in **c**, the mean of the modal peak across all cell densities was between 129 and 152 nm. The plotted curves in **b** and the quantification in **c** are the mean of three independent experiments. **d-e**, TEM micrographs of EVs obtained from Opti-MEM conditioned by SUM159 cells for 24 hours. EVs were enriched by ultracentrifugation (110,000 x g for 1 hour) and the resulting pellets were negatively stained with uranyl acetate for visualization. A magnification of the area delimited by the black dashed square in **d** is shown in **e. f**, Illustration of the proposed relationship between cell density and EV retrieval from the supernatant. When cells are sparsely distributed, EVs can be released from both the apical and the lateral surfaces of the cells and promptly reach the extracellular space (top panel). In contrast, at high cell density only EVs released from the apical surface can promptly reach the extracellular space (bottom panel).

Nevertheless, the experiments presented so far could not discriminate between alternatives: increase in cell density can either prevent EV release, or promote EV internalization and degradation in neighboring cells^18–21^. To test this, we developed an unbiased approach to quantify the fraction of cells that accumulate EVs. The experiment was based on a co-culture of EV-donor and EV-recipient cells. EV-donor cells were transfected with the tetraspanin CD9 conjugated at its N-terminal with the monomeric fluorescent protein Emerald (CD9-Emerald), an improved version of EGFP^22^. CD9-Emerald distributed on the cell surface (Fig. S1d) and its over-expression did not influence EV release either quantitatively or qualitatively (Fig. S1e-g). The EV-recipient cells were transfected with nuclear-tagBFP (n-BFP). Co-culture experiments were set up using both SUM159 and HeLa cells, two models that have been used extensively in EV and membrane trafficking studies^17,23,24^. One day after transfection, respective donor and recipient cells were plated in the same well (Fig. 2a). We reasoned that the inhibition of vacuolar H^+^- ATPase by Bafilomycin A1 (Bafilomycin) could define whether internalized EVs are degraded in lysosomes. For this reason, we added either 50 nM Bafilomycin, a concentration that had non-toxic effects on our cell models (Fig. S2a), or its solvent (DMSO) to the culture dishes. The following day, we resuspended and analyzed the cells by flow cytometry. Cells retaining n-BFP and CD9-Emerald signal accounted for the fraction of n-BFP cells that engulfed CD9-Emerald EVs. In some experiments, the cells were sorted and plated on coverslips to confirm by confocal microscopy that EVs labelled by CD9-Emerald were indeed internalized by the recipient cells (Fig. 2b). The fraction of n-BFP cells that engulfed CD9-Emerald labelled EVs significantly increased in both SUM159 and HeLa cells upon exposure to Bafilomycin (Fig. 2c-e). We confirmed our results by confocal microscopy. In this set of experiments, recipient cells were labelled with clathrin-light chain mRuby (Clathrin-mRuby)^25^ to clearly visualize their surface (Fig. 2f,g). Upon Bafilomycin treatment, CD9-Emerald EVs accumulated in the cytosolic space of the recipient cells (Fig. 2 h,i). An increase in EV release upon Bafilomycin treatment^26^ may have partially contributed to the accumulation of EVs in the endocytic pathway. Thus, to assess whether Bafilomycin enhanced EV release in our cell models, we quantified the number of EVs released by cells cultured in Opti-Mem media in the presence of the drug or its solvent by NTA. We detected ~3 times more EVs upon Bafilomycin administration (Figure S2b). When the supernatants of cells treated or not with Bafilomycin were probed for CD63 and CD9 by slot blot (Fig. S2c), we observed that while both CD63 and CD9 signals increased, the former was significantly more than the latter (Fig S2d). A possible explanation is that Bafilomycin promotes the accumulation of intraluminal vesicles by impairing lysosome function, thereby enhancing the release of exosomes, which are typically enriched in CD63^27^. Alternatively, the increase of CD9^+^ EVs in the supernatant of cells exposed to Bafilomycin may be due to the increased recycling of internalized ectosomes that would otherwise be degraded. Since both the total number of EVs and the fraction of cells that engulfed EVs increased upon Bafilomycin treatment, we concluded that EV release is not hindered by cell-to-cell contact. Instead, increasing cell density results in more EVs being internalized at the cell-cell interface and degraded in the endo-lysosomal network of recipient cells.

**Figure 2.**
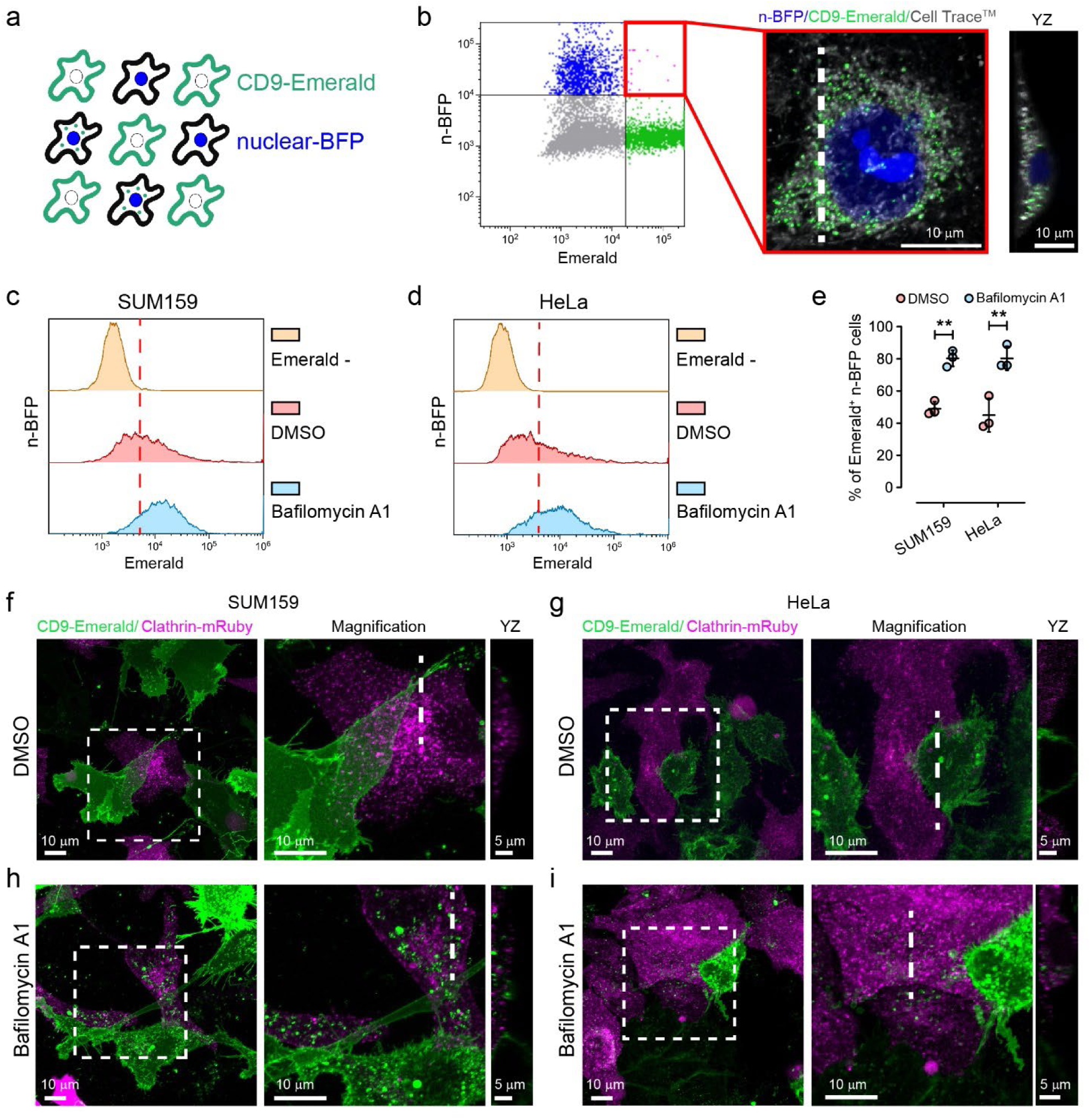
Endosomal alkalization reveals EV exchange in high-density cultures. **a-e**, Flow cytometry assay to define the fraction of recipient cells that engulf EVs. **a**, Scheme representing the co-culture of CD9-Emerald and n-BFP cells. **b**, representative flow cytometry analysis (left) and confocal microscopy analysis of sorted n-BFP^+^/CD9-Emerald^+^ HeLa cells (right). The confocal Z projection and relative YZ profile (computed at the white dashed line) show that EVs are engulfed in the intracellular space of the recipient cells. The confocal data is composed of 40 planes spaced every 250 nm on the z axis. **c-d**, The fraction of SUM159 and HeLa cells recipient cells (n-BFP^+^) internalizing EVs (n-BFP^+^/CD9-Emerald^+^ cells) was quantified by flow cytometry. The co-cultures exposed to Bafilomycin or DMSO are shown in blue and red respectively. In yellow, cultures solely composed of n-BFP^+^ cells (Emerald-) were used as negative controls to define the threshold for Emerald^+^ signal (red dashed line). **e**, Statistical analysis of three independent co-culture experiments (mean ± SEM). The fraction of cells that engulfed EVs increased from 49 ± 2% to 80 ± 3% (1.6 times; mean ± SD) in SUM159 and from 45 ± 6% to 80 ± 4% (1.8 times; mean ± SD) in HeLa cells when treated with Bafilomycin in comparison to the vehicle. Data were compared using a two-tailed t-test where ^**^ p-value < 0.001. **f-i**, Co-cultures set up using CD9-Emerald donor cells and Clathrin-mRuby recipient cells were imaged by live cell confocal microscopy. Donor and recipient cells were mixed and seeded at a 1:1 ratio to reach 80% confluence after 48 hours. Experiments were performed using either SUM159 (**f**,**h**) or HeLa cells (**g**,**i**). After 24 hours of incubation, the co-cultures were exposed to 50 nM Bafilomycin A1 or its solvent (DMSO) for an additional 24 hours. Overall, these experiments suggest that upon internalization, EVs are trafficked through the cell and degraded in lysosomes.

We reasoned that co-culture experiments could be extended to the visualization of EV exchange in real time. To this end, we used lattice light sheet microscopy (LLSM) since image acquisition is faster and less phototoxic than with confocal microscopes. LLSM permits visualization of the entire cell volume while retaining single object/single molecule resolution, resulting in one of the most effective approaches to study cell biology events in five dimensions^24,28,29^. SUM159 donor cells were transfected with CD9-Halo and labelled with 5 nM of photostable JF549-Halo dye (Fig. S3a)^24,30^. As expected, CD9-Halo labelled the cell surface including filopodia and membrane ruffles (Fig. S3b). In addition, when SUM159-CD9-Halo cells were co-cultured with cells expressing CD63-EGFP, CD9-Halo-EVs accumulated in endo-lysosomal structures labelled by CD63 (Fig S3c), recapitulating our previous findings. We used SUM159 cells gene-edited to express the subunit *mu* of the adaptor protein complex AP2 labelled with EGFP (AP2M1-EGFP), a well-established marker of clathrin mediated endocytosis^31^, as recipient cells. While it is well known that clathrin coated vesicle formation takes roughtly 60 seconds and that a frequency of volume acquisition of 0.3 Hz is sufficient to detect the dynamics of clathrin mediated endocytosis^24,31^, the time course of EV release and uptake in real time is poorly investigated. Since a particle of 100 nm in diameter moves at ~10 μm^2^/s when freely diffusing in medium^32^, we reasoned that to successfully track single EVs from release to uptake, it was necessary to focus on the cell-cell interface. At this location, EVs entrapped between contiguous cells would be subjected to a constrained diffusion. Although we collected 23 movies ranging from 15 to 40 minutes of acquisition (a total of 12 hours of imaging), we only identified 10 events that unambiguously corresponded to EVs released from donor cells that were internalized in recipient cells (Fig 3a,b and 3d,e; Movie 1 and 2). At least two reasons explain why the EV internalizations were rarely detected: first, because a cell releases roughly 3,400 EVs in 24 hours (see Fig 1a), which corresponds to only ~3 EVs per minute; second, because we were restricting our studies to a subfraction of EVs by solely monitoring vesicles labelled with CD9. Despite these limitations, we could observe EV exchange in real time. EVs either budded from the cell surface or were released upon the sudden rupture of the tips of filopodia, strengthening the general assumption that CD9 is a marker of ectosomes. In addition, filopodia can undergo additional scission events after being released, producing smaller EVs (Movie 3). Noteworthy, the few EVs that were internalized did not colocalize with AP2-EGFP at any point in time, while other CD9-EVs that were entrapped in clathrin coated vesicles were never endocytosed (Fig. 3c). These results suggested that CD9-EVs are not internalized by clathrin-mediated endocytosis. To unequivocally confirm our observations, we used a HeLa cell model in which complete depletion of clathrin heavy chain could be obtained by activating a doxycycline-dependent shRNA system (Fig. 3f)^33^, thereby inhibiting clathrin-mediated endocytosis (Fig. 3g). HeLa cells depleted or not of clathrin were transfected with n-BFP and co-cultured with CD9-Emerald HeLa cells. Flow cytomtery analysis showed that clathrin depletion did not impair the endocytosis of CD9^+^ EVs (Fig. 3h,i), strengthening our conclusions.

**Figure 3.**
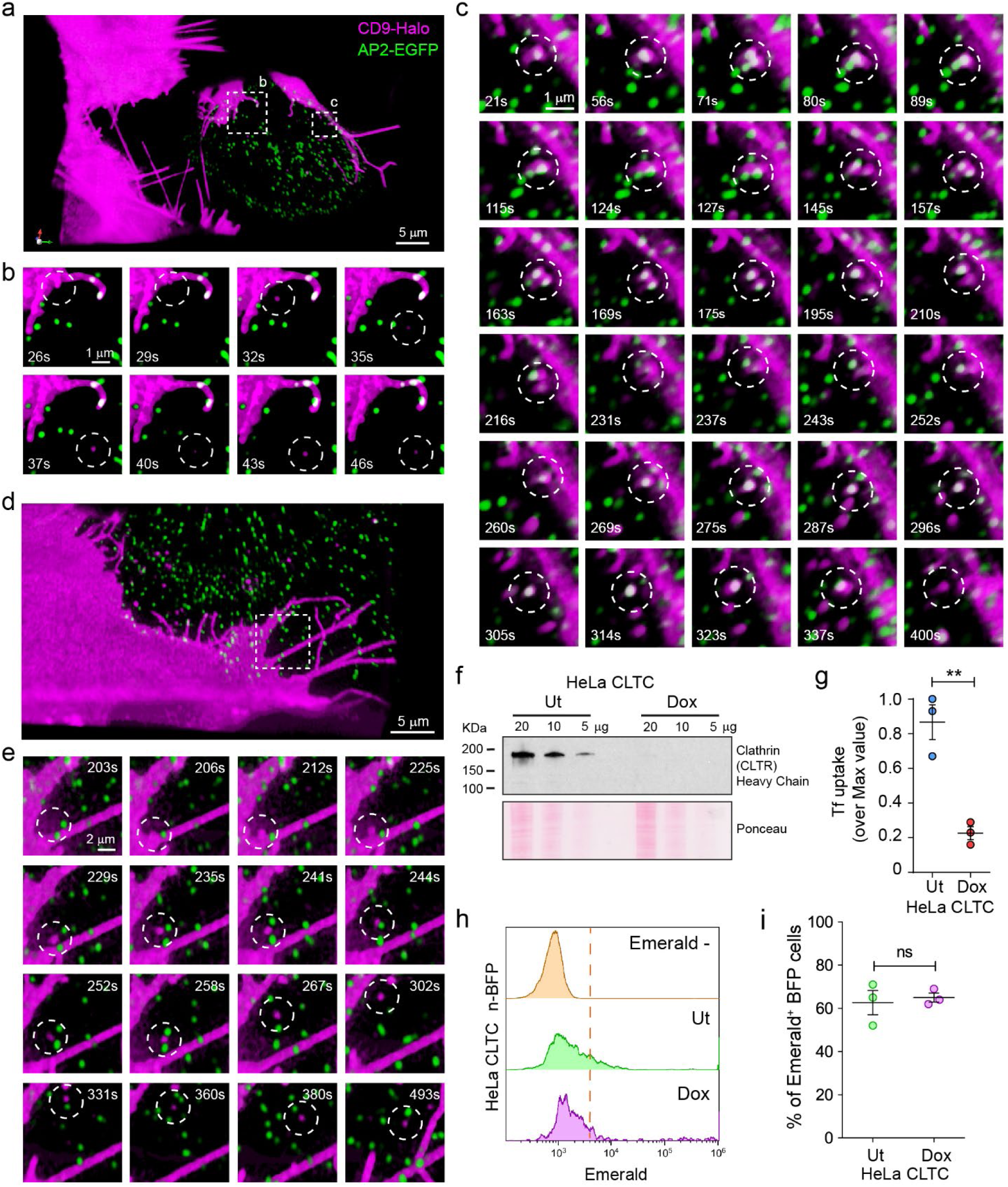
5-D microscopy reveals both the dynamics of EV transfer, and that EV uptake is independent of clathrin-mediated endocytosis. **a-e**, SUM159 cells expressing either CD9-Halo (EV donor) or AP2-EGFP were mixed and seeded at a 1:1 ratio, stained with 5 nM JF549 for 1 hour, and imaged by Lattice Light Sheet Microscopy (LLSM). All movies were acquired in sample scan mode and were composed of sequential volumes of 73 × 53 × 13 μm (corresponding to 40 sequential optical sections acquired with 35 ms of exposure for each channel and spaced every 350 nm after de-skewing; the final frequency of acquisition of the movie was of 0.3 Hz). **a**,**d**, Three-dimensional renderings extrapolated from Movie 1 and 2. The dotted squares define the regions corresponding to the magnified image sequences plotted in panels b, c, and e. **b**,**e**, Internalized CD9^+^ EVs. The lack of co-localization between CD9-Halo and AP2-EGFP upon vesicle internalization (white circle) suggests that clathrin mediated endocytosis (CME) does not support CD9^+^ EV uptake. **c**, An example of a CD9^+^ EV that remained at the cell-cell interface for more than 5 minutes without being efficiently internalized by clathrin coated vesicles despite recurrent colocalization with AP2-EGFP puncta (see also Movie 1). **f-i** To assess the role of CME in CD9^+^ EV uptake, HeLa cells stably expressing a doxycycline (Dox)-inducible shRNA sequence against clathrin heavy chain (HeLa CLTC) were cultured in the presence of Dox for one week to downregulate clathrin expression. **f**, Immunoblotting demonstrated that the Dox-inducible shRNA system abolished the expression of clathrin heavy chain as compared to untreated (Ut) cells. **g**, Clathrin heavy chain depletion impairs CME. HeLa CLTC depleted (Dox) or not (Ut) of clathrin heavy chain were exposed to fluorescently labelled transferrin (A647-Tf), a CME cargo, for 15 minutes at 37°C to allow the ligand to internalize (Tf_INT_), or at 4°C to label the surface receptor by preventing its endocytosis (Tf_SRF_). After washes, the cells were analyzed by flow cytometry. The graph shows the ratios between Tf_INT_ and Tf_SRF_ represented as % of the maximal value (three independent experiments, mean ± SEM). Data were compared using a two-tailed t-test. ^**^ p-value < 0.001. **h**, HeLa cells expressing CD9-Emerald were co-cultured at high density (80% confluence at 48 hours) with either HeLa CLTC Ut-n-BFP or HeLa CLTC Dox-n-BFP. 24 hours post-seeding, the cells were treated with 50 nM Bafilomycin for 4 hours before quantifying the fraction of double-positive CD9-Emerald/n-BFP cells by flow cytometry. The results from three independent experiments were summarized in **i** (data analysis was performed using a two-tailed t-test; ns = not significant) and indicated that CME does not significantly contribute to the uptake of CD9^+^ EVs.

If it is true that EVs are mainly exchanged at the cell-cell interface, high cell density may maximize the number of cells that internalize EVs. To explore this possibility, we seeded co-cultures of HeLa cells expressing either CD9-Emerald (EV-donor) or n-BFP (EV-recipient) at decreasing density. Cells were also exposed to Bafilomycin (as described previously) to maximize the detection of EV exchange. Then, each co-culture was analyzed by flow cytometry to assess the fraction of n-BFP cells that internalized EVs. As expected, cells at confluence had the largest amount of double-positive cells while the fraction was significantly reduced at lower densities (Fig. 4a,b). Since low cell density may result in an inefficient EV exchange due to the decrease in the absolute number of the EVs released in the extracellular space, we set up a compartmentalized co-culture system where the same number of EV donor and recipient cells were seeded in either a *Close* or *Far* configuration. To achieve this, we printed *ad hoc* cell culture rings in a biocompatible material using a stereolithography 3D printer (Fig. S4a,b). The rings were used to plate the cells in a confined space (Fig. 4c; S4c). After cell spreading occurred, the rings were removed and media containing Bafilomycin was added to fill the dish (Fig. S4d). The following day, the cells were detached and analyzed by flow cytometry to quantify the amount of EV transfer. When cells were plated in the *Close* configuration, the fraction of double-positive cells was significantly higher in comparison with the cells plated in the *Far* configuration (Fig. 4d,e). These results revealed that cell-to-cell proximity has a significant impact on the amount of intercellular transfer of EVs, further confirming that EVs are exchanged locally.

**Figure 4.**
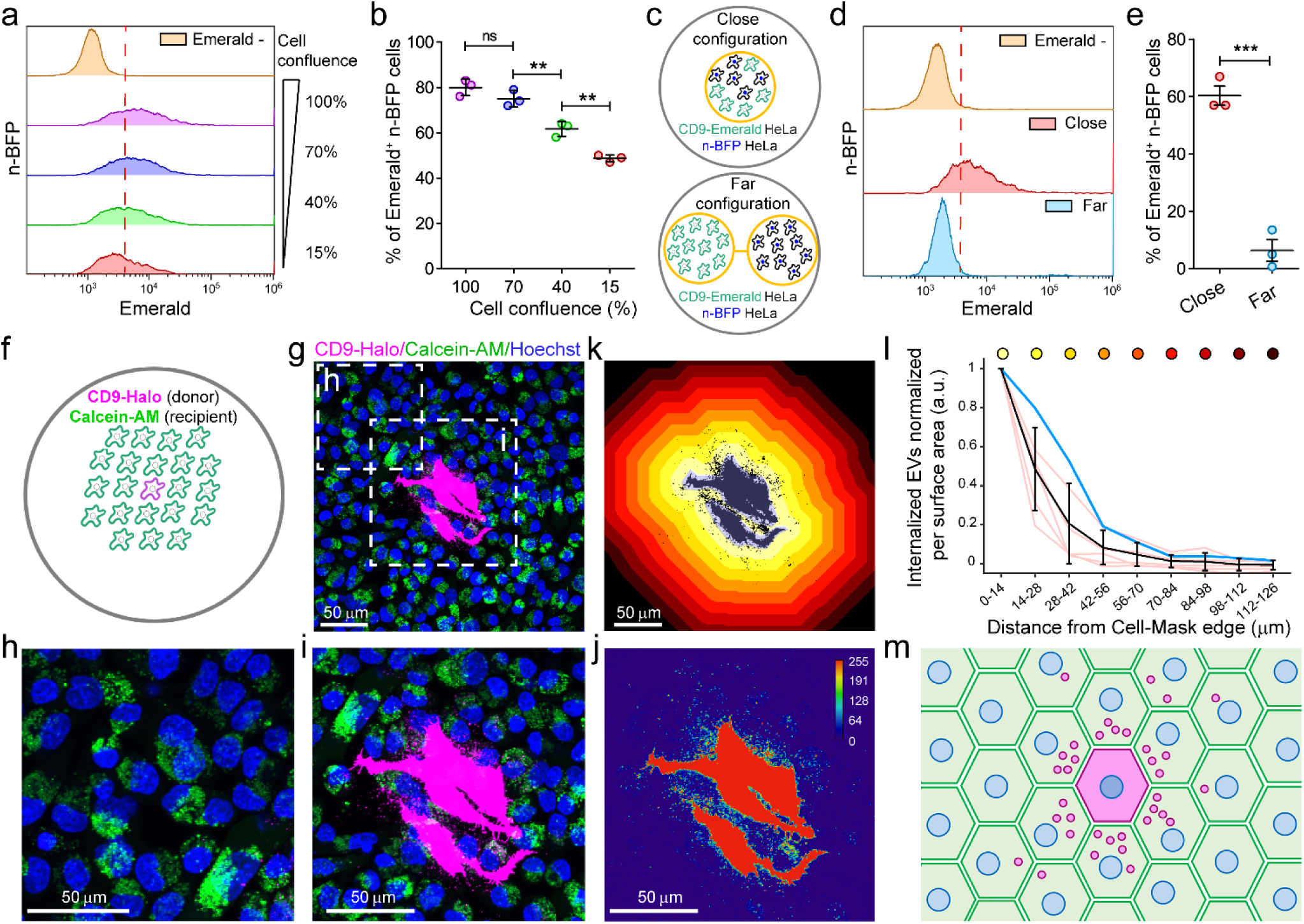
EV exchange is favored by cell proximity. **a**,**b** Co-cultures of HeLa cells expressing either n-BFP or CD9-Emerald were used to assess the impact of cell proximity on the efficiency of EV exchange. To do so, EV donor (CD9-Emerald) and EV recipient (n-BFP) cells were seeded at decreasing cell densities (100%-15%; confluence at 48 hours). 24 hours post-seeding, the cells were treated with 50 nM Bafilomycin for an additional 24 hours before analysis by flow cytometry. **a**, The density plots show the density distributions of n-BFP^+^ cells according to the CD9-Emerald signal, in function of the cell density. The double-positive cell populations correspond to the events in the histograms on the right side of the red line, which marks the threshold for Emerald^+^ signal, defined using the control population (nBFP^+^/Emerald^-^ cells). The results from three independent experiments (mean ± SD) are shown in the graph in **b**. Data were compared using One-way ANOVA and Tukey’s *post-hoc* test (ns = not significant; ^**^ p-value < 0.001). **c-e**, To further explore the effect of the intercellular distance on the amount of EV transfer, custom-made rings (1 cm radius, 0.5 cm height) were printed in a biocompatible material using a stereolithographic 3D device (Formlabs). The rings were used as mobile chambers to control the distance between EV donor and EV recipient cells when plating. The pictogram in **c** outlines the *Close* and *Far* configurations that characterize the experiment (for more details, see Fig S4 a-d). *Close* configuration (Top panel): EV donor (CD9-Emerald) and EV recipient cells (n-BFP) were seeded together at a 1:1 ratio in one ring in the middle of the dish (100% final confluence) to generate a dense co-culture. *Far* configuration (Bottom panel): using two connected rings (4 cm apart), an equal number of EV donor (CD9-Emerald) and EV recipient (n-BFP) cells were seeded in each ring. After 24 hours, the rings were removed in both conditions and the dishes were filled with complete media supplemented with 50 nM Bafilomycin. 24 hours post-treatment, the cells were detached and analyzed by flow cytometry to compare the efficiency of EV transfer when the two cell types were seeded together versus separately. One representative experiment is shown in **d** while data from three independent experiments are summarized in **e. d**, n-BFP^+^ cells seeded without donor cells were used as negative controls to define the threshold for Emerald^+^ signal (yellow distribution and red dashed line). The red and blue distributions depict the CD9-Emerald intensities that were recorded in the n-BFP^+^ recipient cell population when co-cultured in either the *Close* (red) or *Far* (blue) configurations with CD9-Emerald donor cells. **e**, When cells were cultured in the *Close* configuration, 60 ± 6% (mean ± SEM) of n-BFP^+^ cells had internalized CD9-Emerald^+^ EVs. In contrast, only 6.4 ± 6% (mean ± SEM) of n-BFP^+^ cells cultured in the *Far* configuration received EVs from the CD9-Emerald donor cells located 4 cm away. The differences in EV transfer between the two configurations were statistically significant (data analysis was performed using a two-tailed t-test where ^***^ p-value < 0.0001). **f-i**, To assess the propagation of EVs between adjacent cells, we set up a co-culture system as depicted in **f**. HeLa cells expressing CD9-Halo were co-seeded on glass coverslips with HeLa cells labelled with Calcein-AM at a 1:3,000 ratio and 80% seeding density to maximize the possibility that EV donor cells were surrounded by a confluent monolayer of recipient cells after 48 hours. 24 hours post-seeding, the cells were treated with 50 nM Bafilomycin for 24 hours. Before imaging, the cells were incubated with 5 nM JF549 for 1 hour to label CD9-Halo. Additionally, the cells were incubated with Hoechst for 10 minutes to stain nuclei. After two washes in complete media, the cells were imaged by confocal microscopy (**g**). Magnification of an area at the edge of g (**h**) shows negligible CD9-Halo fluorescence in comparison to the accumulation of CD9^+^ EVs observed within recipient cells in immediate proximity to the donor cells (**i, j**). **k**, To quantify the accumulation of EVs in recipient cells within the context of their proximity to donor cells, we developed a segmentation analysis that correlates CD9-Halo fluorescence intensity with distance from the edge of the nearest donor cell. We did this by normalizing the absolute CD9-Halo intensity to the respective segmented surface area. The results in **l** show that the majority of EVs are transferred to cells that are within 28 μm of the edge of a donor cell. The color-coded dots correspond to the segmented surface areas shown in **k. m**, Illustration of the proposed mechanism of EV transfer. Cell density is a major determinant of EV exchange. When cells are densely packed, EVs are captured by adjacent cells and only a limited fraction can reach extracellular fluids. Moreover, fast diffusion of EVs in extracellular space limits their capture by distant cells. These data, when extrapolated to three dimensions, suggest that EVs detected in bodily fluids mainly originate from cells directly exposed to them. Consequently, blood and endothelial cells are the main source of circulating EVs while the contribution of other tissues is limited. However, changes in physical parameters such as intraparenchymal pressure and endothelial permeability may promote EV escape. Since these properties are strongly influenced by pathological conditions such as inflammation, these data provide experimental evidence to support the notion that EVs are important resources for the development of liquid biopsy approaches.

However, the motion of bodily fluids may propel EVs to reach anatomical districts far from the site of origin. To simulate this event, CD9-Emerald and n-BFP^+^ HeLa cells seeded in the *Far* configuration were incubated either motionless or under continuous agitation in the presence of Bafilomycin. A co-culture of CD9-Emerald and n-BFP HeLa cells plated in the *Close* configuration was used as a reference (Fig. 4c; S4c). Interestingly, when cells were analyzed by flow cytometry, there was no significant difference in the fraction of cells that internalized EVs between the two conditions when seeded in the *Far* configuration (Fig. S4e,f). Conversely, the fraction of internalized EVs in the cells plated in the *Close* configuration was significantly higher when compared with both samples plated in the *Far* configuration (Fig. S4f). These results confirmed that EVs are mainly exchanged at the cell-cell interface. These observations raise an additional question: how far can EVs travel between adjacent cells? To explore this, we plated CD9-Halo HeLa cells with HeLa cells loaded with Calcein AM in a ratio of 1:3,000 to ensure that donor cells would be surrounded by a monolayer of recipient cells (Fig. 4f). After 48 hours, the cells were imaged by confocal microscopy to capture images of the sparse donor cells (Fig 4g-j). We developed a segmentation protocol that identifies the donor cells and designs concentric regions of interest to measure the relative intensity of EV signal in function of the distance from the edges of the donor cells (Fig. 4k). Our results demonstrate that EV signal decays exponentially from the contours of the donor cell (Fig. 4l) with the maximum being within the first 14 μm, which roughly corresponds to the first layer of calcein-labelled recipient cells.

In conclusion, our results support a model where EVs are exchanged between adjacent cells and promptly degraded, suggesting that EVs are mainly confined within the tissue of origin (Fig. 4m). It may seem difficult to reconcile this observation with the role that EVs play as mediators of intercellular communication. However, it is important to consider that many of the effects that EVs elicit are short range communication in nature, such as the trophic support that EVs released by Schwann or glial cells can provide to damaged axons^34^, or EVs exchanged at the immune synapse^35^. Moreover, this communication can be regulated by ligands harbored on the EV surface that can stimulate cells by clustering signal receptors^36,37^ and therefore act at the cell surface before being degraded in lysosomes. Although we did not detect EV fusion with recipient cells in our system, it is an event that can happen, albeit rarely ^23^. Finally, when extrapolated to three dimensional tissues, our data predict that circulating EVs originate from cells exposed to the bloodstream. These could be endothelial and blood cells, or tissues with leaky vasculature such as the liver. In addition, our data predict that EVs can escape from tissues when permeability and intraparenchymal pressure increase. Since these are the typical events that characterize inflammation, which is a hallmark of chronic diseases including cancer, our data provide physiological evidence to support the use of EVs as effective and specific tools for the development of liquid biopsy approaches.

## MATERIALS AND METHODS

### Reagents

The human triple-negative breast cancer cell line SUM-159PT and the gene edited cell line AP2S1-EGFP (SUM-AP2-EGFP) were a kind gift of Dr. Tomas Kirchhausen (Harvard Medical School, Boston, MA)^31^; the human cervical cancer cell line HeLa and inducible knock down for clathrin (HeLa CLTC) were from Drs. Pier Paolo Di Fiore and Sara Sigismund (IEO, European Institute of Oncology, Milano)^33^. The mammalian expression vectors for Clathrin-mRuby, mEmerald-CD9, and mTagBFP-Nucleus-7 were generously provided by Dr. Michael Davidson (Addgene Plasmids: #55852, #54029 and #55265), while the one to express EGFP-CD63 was provided by Dr. Martin Hemler. JF549-Halo Tag ligand is a kind gift of Dr. Luke Lavis (Janeila Research Campus). The expression vectors were propagated in DH5-alpha *E. coli* (New England Biolabs) and plasmids purified using Plasmid Miniprep or Midiprep Kits (Qiagen) or a PureLink HiPure Plasmid Filter Midiprep Kit (Thermo Fisher Scientific). Primers were obtained from IDT (Integrated DNA Technologies). Primary antibodies against clathrin heavy chain (mouse monoclonal, 610499, BD Biosciences), CD9 (mouse monoclonal, 2897163, EMD Millipore), and CD63 (mouse monoclonal 561925, eBiosciences) were used for Western blotting together with anti-mouse IgG antibody conjugated with horseradish peroxidase (goat-polyclonal, 5178-2504, Bio-Rad). Chemicals were purchased from Sigma-Aldrich unless specified otherwise.

### Cell handling and transfection

Cell lines were cultured at 37°C and 5% CO_2_; SUM159 were grown in F12-Glutamax (Thermo Fisher Scientific) supplemented with 5% fetal bovine serum (FBS) (VWR), 100U/ml penicillin and streptomycin (Thermo Fisher Scientific), 1 μg/ml hydrocortisone (H-4001; Sigma-Aldrich), 5 μg/ml insulin (128-100; Cell Applications), and 10 mM 4-(2-hydroxyethyl)-1-piperazine-ethane-sulfonic acid (HEPES), pH 7.4. HeLa cells were grown in DMEM-high glucose supplemented with 10% FBS, 100U/ml penicillin and streptomycin, and 1% Glutamax (Gibco). To induce clathrin knock down, HeLa CLTC cells were cultured for 1 week in DMEM-high glucose supplemented with 10% Tetracycline-free FBS (Thermo Fisher), 100U/ml penicillin and streptomycin, 1% Glutamax (Gibco), and 0.5 μg/ml doxycycline (Sigma-Aldrich). Cells were resuspended to be propagated or subjected to other experimental procedures by exposure to 0.25% trypsin-EDTA (Thermo Fisher Scientific) for 3 minutes at 37°C, followed by a wash in 10 ml complete media and centrifugation at 1,000 rpm for 5 minutes. Cells were counted and plated as needed. Cell culture flasks and multi well plates were purchased from Greiner Bio-One.

For transient transfections, 4 × 10^5^ cells were plated in one well of a 6-well plate in complete medium. The following day 3 µl of Mirus TransIT 2020 reagent (Mirus Bio, Madison, WI) and 1 µg of plasmid were combined in 250 µl Opti-MEM (Thermo Fisher Scientific). After 30 minutes the mixture was added dropwise to the cells. Expression of Clathrin-mRuby, CD9-Emerald, nuclear-BFP, CD9-Halo, and CD63-GFP were obtained by transient transfection. To obtain SUM159 cells stably expressing CD9-Halo, the transiently transfected cells were expanded in T75 flasks in the presence of the antibiotic G418 (800 μg/ml for 15 days), and further enriched by two rounds of cell sorting and expansion over the course of three weeks.

### Plasmid assembly

The mammalian expression vector encoding for CD9-Halo was developed starting from the mEmerald-CD9 vector and a previously developed CALM-Halo construct^24^. Emerald was substituted with HaloTag using an approach based on isothermal assembly^38^. Briefly, two overlapping PCR products were generated using the high-fidelity DNA polymerase Q5 (New England Biolabs): one was the linearized recipient CD9 plasmid; the second was the HaloTag insert. Primers to generate the CD9 linearized plasmid were:

FW: 5’-cgagatttccggcggcagcggagggCCGGTCAAAGGAGGCACC-3’ RW: 5’-tttctgccatGGTGGCGACCGGTAGCGC-3’

Primers to amplify the HaloTag insert were:

FW: 5’-ggtcgccaccATGGCAGAAATCGGTACTG-3’

RW: 5’-ctttgaccggccctccgctgccgccGGAAATCTCGAGCGTCGAC-3’

Upper-case characters in the primer sequences indicate the regions that anneal with the original constructs. The lower-case characters indicate additional bases inserted to provide matching sequences between the two amplified PCR products and a flexible linker (SGGSGG) between the HaloTag and the CD9 sequence. The PCR products were ligated using the NEBuilder HiFi DNA Assembly kit (New England Biolabs, Ipswich, MA) and transformed into competent DH5a *E. coli* (5-alpha Competent E. coli) following the protocol provided by the company (New England Biolabs, Ipswich, MA). For colony selection, bacteria were plated on Luria Broth agar plates supplemented with kanamycin (25μg/ml). Plasmids were extracted using the QIAprep Miniprep Kit (Qiagen, Hilden, Germany) or HiPure Plasmid Filter Midiprep Kit (Thermo Fisher Scientific, Waltham, MA) and sequenced to ensure that the correct modifications were in place.

### Cell protein extraction and immunoblotting

HeLa and HeLa CLTC cells grown on 6 well plates at 80% confluence were detached, washed three times in phosphate-buffered saline (PBS), and solubilized at 4°C in lysis buffer (50 mM Hepes pH 7.4, 150 mM NaCl, 15 mM MgCl2, 1 mM EGTA, 10% glycerol, and 1% Triton X-100) supplemented with a protease inhibitor mixture (Roche). Nuclei were removed from the lysates by centrifugation (800 x g, 10 minutes, 4 °C). Protein concentration was measured by Bicinchoninic acid (BCA) assay (Pierce). Samples were loaded in precast 4-20% gradient sodium dodecyl sulfate (SDS) polyacrylamide gels (Mini-PROTEAN TGX gels, Bio-Rad) and run at room temperature with 100 V (constant) before being transferred to polyvinylidene fluoride or nitrocellulose membranes using the Trans-Blot Turbo system (Bio-Rad) for 10 minutes at 2.5 A. Membranes were stained with Ponceau S solution then saturated in blocking buffer composed of Tris Buffer Saline (TBS) (150 mM NaCl and 50 mM Tris pH 7.4) supplemented with 5% nonfat dry milk and 0.05% Tween20. The membranes were then incubated for 1 hour with the appropriate primary antibodies diluted in blocking buffer (100 ng/ml). After three washes of 5 minutes apiece in blocking buffer, the membranes were incubated with 1 μg/ml of secondary antibody for 1 hour. After extensive washes in Tris-buffered saline, the chemiluminescent signal was elicited by 5 minute incubation with enhanced chemiluminescence substrate (ECL, Pierce, Thermo Fisher Scientific) and captured photographically on film (CL-XPosure, Thermo Fisher Scientific).

### Slot blot for EV protein detection

SUM159PT cells were seeded in 6 well plates at increasing density. 24 hours post-plating, the cells were washed twice with PBS and maintained in Opti-MEM (1 ml/well). After 24 hours, the cell supernatants were collected and centrifuged at 5,000 x g for 10 minutes to remove floating cells and debris. The resulting supernatants were spotted on a nitrocellulose membrane (0.45 um, Bio-Rad) using a slot blot apparatus. Briefly, pre-wet nitrocellulose in TBS solution (150 mM NaCl, 50 mM Tris pH 7.4) was inserted into a slot blot apparatus (Bio-Rad) connected to vacuum aspiration. The samples (200 µl each) were loaded into the slot-blot wells and dried out by vacuum. The process was repeated until the entire volume of each sample was loaded (1 ml each sample). An equal final volume of Opti-MEM was spotted on separate wells as negative control following the same procedure.

The nitrocellulose membrane was dried at RT for 2 hours, then saturated for 1 hour with blocking buffer (5% non-fat dry milk in TBS + 0.05% Tween20). The membrane was incubated overnight at 4°C with the primary antibodies diluted in blocking buffer (100 ng/ml). The day after, the membrane was washed three times with blocking buffer, incubated with anti-HRP secondary antibodies, and the signal developed with ECL reaction. Densitometric analysis were performed with ImageJ software (NIH).

### Nanoparticle Tracking Analysis (NTA)

Cells were seeded in 6 well plates and grown in complete media. The day after cells were washed four times with HBSS (Gibco) and incubated in Opti-MEM (Gibco). After 24 hours, the cell supernatants were collected and centrifuged at 5,000 x g for 10 minutes to remove floating cells and debris. The resulting supernatants were collected and kept on ice before injection into the imaging chamber of NanoSight NS300 (Malvern Panalytical) by a syringe pump. Three videos of at least 30 seconds each were recorded for each sample. The instrument is equipped with a cMOS camera and controlled by NTA software (version 3.3). Camera shutter speed was set to 25 fps, camera gain to 250, and detection threshold to 5. Minimum expected particle size and minimum track length were set to automatic.

### Live cell confocal microscopy and data analysis

After transfection, cells were detached, counted, and mixed in a Falcon tube. The cell mixture containing EV-donor (CD9-Emerald or CD9-Halo) and EV-recipient cells (Clathrin-mRuby, n-BFP, AP2-EGFP or CD63-GFP) were plated on a 6 well plate containing glass coverslips (#1.5; Warner Instruments, diameter 25 mm) previously cleaned by sonication (Ultrasonic Cleaner Branson 1800, Branson) in ethanol and dried at 120°C. For SUM159 co-culture experiments, a total of 3 × 10^5^ cells were seeded per well, while for HeLa co-cultures 6 × 10^5^ cells per well were used. The day after seeding, cells were treated with 50 nM Bafilomycin A1 or its solvent (DMSO) for 24 hours. When CD9-Halo^+^ cells were used as EV donors, the co-cultures were incubated with JF549-HaloTag ligand^39^ (5 nM) dissolved in complete media. After one hour of washes in cell media, the coverslips were moved into a metal chamber for live-cell imaging (Thermo Fisher Scientific) and acquired with a Nikon A1R scanning confocal microscope equipped with a temperature and CO_2_-controlled chamber. The microscope is controlled by the NIS Element software (Nikon). Image stacks were acquired using a Plan Apo 100X oil immersion lens (NA:1.45); planes were spaced 0.25 μm along the z axis and pixel size was 0.25 μm. ImageJ (NIH) was used to project the images across the z axis.

To define the concentration of JF549 Halo ligand that maximized CD9-Halo labelling while preventing nonspecific fluorescent signal, HeLa cells stably transfected with CD9-Halo were plated on 25 mm glass coverslips as described previously, exposed to increasing concentrations of JF549 (2.5-40 nM) for 1 hour, then extensively washed with media for an additional hour. Nuclei were labelled with 2 μg/ml Hoechst (Thermo Fisher Scientific) for 10 minutes. Image stacks were acquired using a Plan Fluor 40X oil immersion lens (NA:1.3); planes were spaced 0.5 μm along the z axis and pixel size was 0.62 μm.

Five fields of view were acquired by live cell confocal microscopy for each coverslip in order to have > 100 cells per condition. The images were analyzed using Cell Profiler^40^, which measured the intensity of each cell in the JF549 channel. A cell was defined by the software as the object comprised within the circle drawn 25 pixels around the perimeter of the nucleus, identified by the Hoechst staining.

To assess the extent of EV exchange at the cell-cell interface, HeLa cells expressing CD9-Halo were seeded on glass coverslips with HeLa cells, labelled for 1 hour with 2 μM Calcein-AM (Thermo Fisher Scientific), at a 1:3,000 ratio (80% confluency at seeding) to maximize the possibility that EV donor cells were surrounded by a confluent monolayer of recipient cells after 48 hours. The day after seeding, cells were treated with 50 nM Bafilomycin A1 for 24 hours. Before imaging, the cells were incubated with JF549 (5 nM, 1 hour of incubation) to label CD9 and Hoechst (10 minute of incubation) to stain nuclei. After extensive washes in complete media, the cells were imaged with the Nikon A1R confocal microscope using a Plan Fluor 40X oil immersion lens (NA:1.3); planes were spaced 0.3 μm along the z axis and pixel size was 0.62 μm.

Fields were selected with CD9-Halo cells in the center of the image. The images were segmented using a customized routine developed in MATLAB. First, the code generated a mask to outline the CD9-Halo cells (origin of the EVs). Then, an algorithm subdivided the rest of the image in concentric surfaces at increasing distance from the edge of the cell. The lateral size of the concentric masks was constant and approximatively set to half of the diameter of an HeLa cell (14 μm).

### Lattice Light Sheet Microscopy (LLSM) live cell imaging, data acquisition and visualization

We imaged HeLa cells with a lattice light-sheet fluorescence microscope (Intelligent Imaging Innovation). Directly derived from the instrument patented by Dr. Eric Betzig and colleagues^28^, this system utilizes Bessel beam lattice sheet illumination via cylindrical lenses and a high-speed spatial light modulator (SLM) for multicolor imaging, as well as an annular mask array for the formation of various light sheets and galvo mirrors to control lattice movement in x and z. It has cameras in the image and Fourier space to inspect the lattice and annular mask shapes. Assembled on an active vibration isolation table equipped with active and passive system for vibration cancellation (TMC, Ametek), it is equipped with four laser lines for sample excitation (405 nm, 300 mW, Omicron; 488 nm, 300 mW, Coherent; and 560 nm, 500 mW and 642 nm, and 500 mW, MPB Communication) and the required filter sets necessary to efficiently image two color sets at a time (Semrock). Two water immersion objectives control the illumination (28×/0.67 NA) and detection (25×/1.1 NA, expanded to reach a final 62.5× magnification). Two piezo controllers operate the translations of the stage and of the objective. The instrument is also equipped with a temperature-controlled, 90 × 1,500 cm (36″ x 60″) specimen chamber maintained at 37°C. Images are acquired by two high-speed, high-resolution 2K x 2K sCMOS camera with 82% QE (ORCA-Flash4.0 V3, Hamamatsu). A computer workstation (Dell) is equipped with the software SlideBookTM 6 (Intelligent Imaging Innovation) that fully controls the instrument (high-speed synchronization of laser firing, SLM pattern display, galvo movements, and imaging camera readout along with the ability to de-skew and view data).

*Data Acquisition and visualization*. To acquire the images, we illuminated the sample with a dithered light sheet generated in the square lattice configuration^28^ using an excitation NA of 0.550/0.493 and full width at half maximum (FWHM) beam length of 15 μm. SUM159 cells stably expressing CD9-Halo or AP2-EGFP were excited for 35 ms with the 488-nm and 561-nm lasers at a power of 12 mW and 27 mW respectively, measured before the cylindrical lenses (500-mW laser, acousto-optic tunable filter 1 [AOTF 1]). Image stacks were composed of 40 sequential planes of 768 × 512 pixels spaced 0.6 μm in sample scan mode (corresponding to 73 × 53 x13 μm volume spaced 0.35 μm over the z axis after de-skewing). The movies we analyzed were composed of a variable number of stacks (from 234 up to 932) acquired every 3 seconds to generate movies spanning from 11 minutes to 46 minutes. For visualization, the de-skewed stacks (Intelligent Imaging Innovation) were deconvolved using the Lucy-Richardson algorithm (MATLAB, MathWorks) and rendered with a 5D viewer developed in MATLAB environment^41^. The 3D rendered images were exported as .tiff images or movies in .mp4 format.

### Flow cytometry and cell sorting

HeLa or SUM159 cells plated on 6 well plates were transfected with CD9-Emerald or n-BFP. After 24 hours, the cells were detached by 0.25% trypsin-EDTA (Thermo Fisher Scientific), counted, and mixed 1:1 in a Falcon tube. For each condition, a total of 120,000 HeLa cells (60,000 CD9-Emerald and 60,000 n-BFP) were mixed. In the experiments using SUM159, a total of 60,000 cells per condition were used (30,000 CD9-Emerald and 30,000 n-BFP).

Mixtures were seeded in multiwell plates and left to adhere and spread for 48 hours. The day after seeding, cells were treated with 50 nM Bafilomycin A1 for 24 hours. Then, the cell co-cultures were detached by 0.25% trypsin-EDTA, rinsed in complete media, and analyzed by flow cytometry (Gallios, Beckman Coulter). Unstained cells and single color controls (CD9-Emerald^+^ and n-BFP^+^) were used to set photomultiplier voltages. Analyses were performed using Kaluza software (Beckman Coulter). Alternatively, double-positive n-BFP/CD9-Emerald cells were sorted using an ARIA III cell sorter (Beckton Dickinson and Company). Sorted cells were seeded on glass coverslip for cell imaging. After spreading, cells were labelled with 1 μM Cell Trace-Far Red (Thermo Fisher Scientific) in serum-free medium for 20 minutes. After two washes in complete medium, coverslips were inserted into a metal chamber for live-cell confocal imaging with a Nikon A1R microscope. Image stacks were acquired using a Plan Apo 100X oil immersion lens (NA:1.45); planes were spaced 0.25 μm along the z axis and pixel size was 0.25 μm. ImageJ (NIH) was used to project the images across the z axis.

### Cell viability assay

A 3-(4,5-dimethylthiazol-2-yl)-2,5-diphenyltetrazolium bromide (MTT) assay was performed to assess cell viability upon treatment with Bafilomycin A1. HeLa cells plated in 96 well plates (10,000 cells/well) were treated with increasing concentrations of Bafilomycin A1 (0-400 nM) or its solvent (DMSO) for 24 hours. Afterwards, MTT (5 mg/ml) was added to each well and incubated at 37°C for 2 hours. The cell media was removed and replaced with DMSO to dissolve formazan crystals, the MTT metabolite. The plate was kept under agitation for 15 minutes, then the absorbance was measured at 560 nm using a plate reader (BioTek SynergyH1).

### Transmission Electron Microscopy (TEM) of EV pellets

SUM159 cells were seeded at 70% confluence in two T175 flasks. After spreading, cells were washed in HBSS and incubated in Opti-MEM for 24 hours. The conditioned media was collected and centrifuged at 5,000 x g for 10 minutes to remove cell debris. The resulting supernatants were transferred into ultracentrifuge-proof tubes and centrifuged for 1 hour and 30 minutes at 110,000 x g using a swing rotor (SW32, Bechman Coulter). The supernatants were discarded and the pellets were resuspended in 60 μl of HBSS. 20 μl of EV suspension were adsorbed onto 400-mesh formvar/carbon-coated grids for 10 minutes at RT. Adherent EVs were stained with uranyl-acetate and immediately imaged using a Tecnai G2 Biotwin transmission electron microscope (FEI).

### Design and fabrication of plastic rings for cell co-culture

Cell rings were designed using the online software Tinkercad (Autodesk). For the *Close* configuration, we designed rings of 1 cm radius and 0.5 cm height. For the *Far* configuration, we designed two rings (1 cm radius and 0.5 cm height) connected by a stem of 4 cm. Rings were 3D-printed in a biocompatible resin (*Surgical Guide*) using a Form 3B+ printer (Formlabs) following the company’s protocol. After printing, the rings were washed extensively in iso-propanol and cured. The curing procedure consisted of simultaneously exposing the printed parts to heat (60°C) and light (405nm) for 60 minutes in a Form Cure station (Formlabs).

Ring patterns were placed into 10 cm Petri dishes. To plate cells in the *Close* configuration, we mixed HeLa cells transfected with CD9-Emerald or n-BFP at a 1:1 ratio. The *Far* configuration consisted of plating donor and recipient cells in two different rings connected by a 4 cm stem. 60,000 HeLa cells transfected with CD9-Emerald were plated in one ring and 60,000 HeLa cells transfected with n-BFP were plated in the other ring. After 24 hours, the culture media and the rings themselves were removed from the dishes. The Petri dishes were then washed in HBSS and the cells were incubated in complete DMEM media. The following day, the cells were treated with 50 nM Bafilomycin A1 for 24 hours. Some plates were placed on an orbital shaker at 37°C to promote media movement while other plates were kept still in the incubator. After 24 hours, cells were detached by trypsinization, centrifuged, and analyzed by flow cytometry to measure the percentage of double-positive CD9-Emerald/n-BFP cells.

### Transferrin-647 uptake to evaluate the functionality of clathrin-mediated endocytosis

HeLa CLTC were treated or not with 0.5 μg/ml doxycylin for one week to induce the expression of shRNA specific for clathrin heavy chain. Then, the cells were washed with HBSS and exposed to 0.5 μg/ml Transferrin-Alexa Fluor 647 (A647-Tf, Thermo Fisher Scientific) dissolved in complete DMEM for 10 minutes at 37°C. Afterwards, the dishes containing the cells were set on ice to block endocytosis and the cells were washed with ice-cold HBSS. To measure the amount of transferrin receptor present at rest on the cell surface, cells in other wells were exposed to A647-Tf at 4°C for 10 minutes. To remove the surface-bound fluorescent ligand, some wells were exposed twice to an acidic solution for 1 minute (150 mM NaCl and 0.1 M glycine, pH 2.5). After two additional washes in ice cold HBSS, the cells were detached by trypsinization, collected in flow tubes containing 1 ml of media to inactivate trypsin, centrifuged at 300 x g, and resuspended in ice-cold media. The samples were then quantified by flow cytometry (Gallios, Beckman Coulter) and the resulting data were analyzed by Kaluza software (Beckman Coulter). Transferrin internalization was normalized to the amount of surface bound. The experiment was repeated three times and the normalized means obtained across the experiments were compared using an unpaired t-test (GraphPad Prism).

### Statistical analysis and figure preparation

Statistical analyses were performed using the software Prism (GraphPad); figures were assembled in Adobe Illustrator (Adobe).

## ACKNOWLEDGEMENTS

We thank Jeff Tonniges (The Ohio State University, CMIF) for helping with TEM sample staining and image acquisition. We acknowledge the use of Microscopy, Genomics, and Flow Cytometry Shared Resources at The Ohio State University, which are supported by the National Cancer Institute (NCI, P30CA016058). We thank the Advanced Imaging Center (AIC) at Janelia Research Campus for training on their LLSFM, John M Heddleston, Eric Wait, Satya Khuon, and Teng-Leong Chew from the AIC team. The AIC is jointly supported by the Howard Hughes Medical Institute and the Gordon and Betty Moore Foundation. E.C. was supported by the OSU Comprehensive Cancer Center and the Ohio Cancer Research via the McCurdy/Kimball Midwest Research Fund. F. C. was supported by an American-Italian Cancer Foundation Post-Doctoral Research Fellowship and by a Pelotonia Post-Doctoral Fellowship.

## AUTHORS’ CONTRIBUTIONS

E.C. conceived and supervised the research; F.C. and E.C. designed research; F.C. and E.G.N. performed research; F.C. and E.C. analyzed data; E.C. wrote the first draft of the manuscript; All authors contributed and agreed on the final version of the paper.

## FIGURES, FIGURE LEGENDS, AND MOVIE LEGENDS

**Figure S1.**
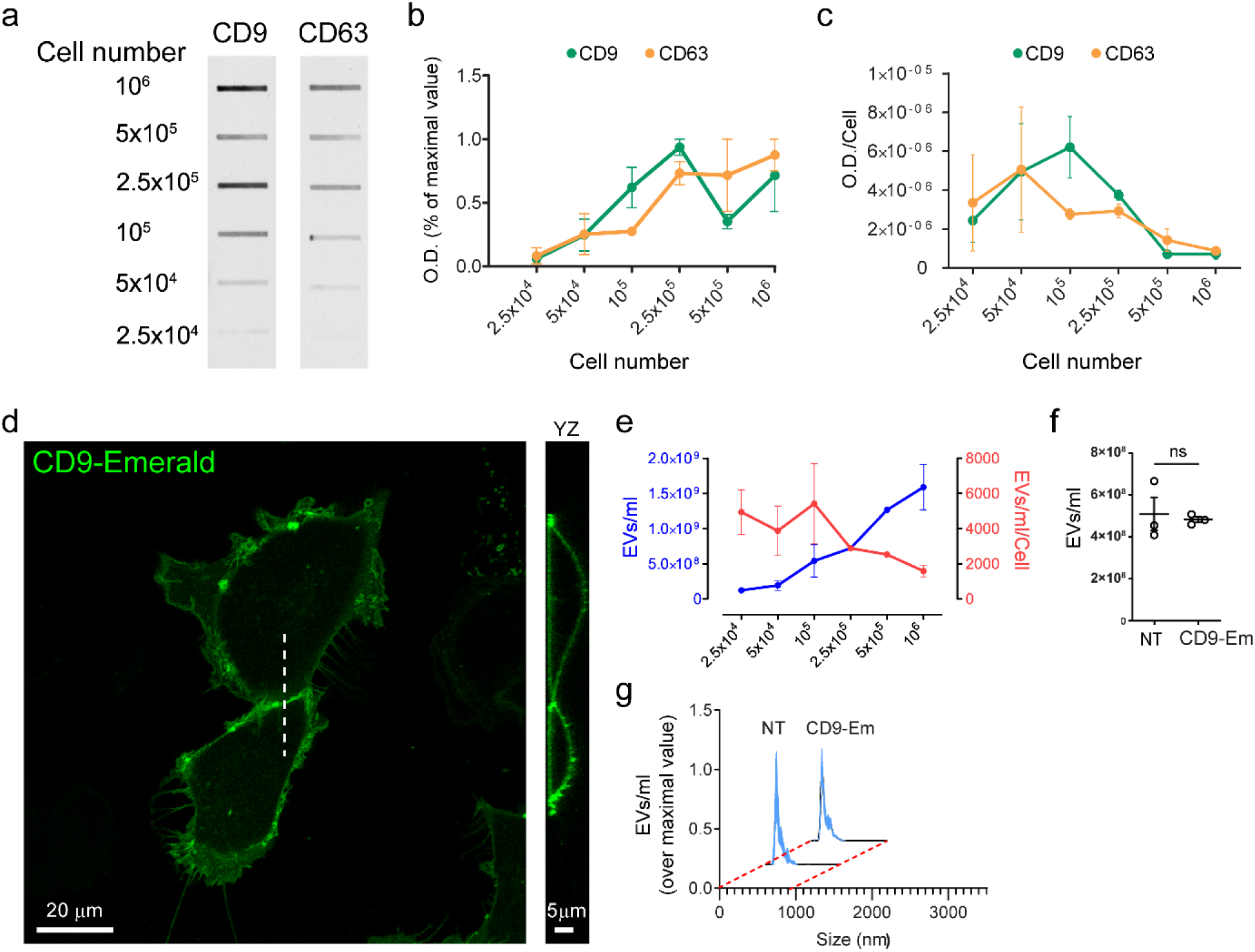
Cell density influences the release of CD9^+^ and CD63^+^ EVs equally and CD9-Emerald overexpression does not affect either the number or the size distribution of EVs. **a-c**, Cell supernatants, from the experiment in Fig.1, cleared from debris were immunoblotted to assess the relative amount of ectosomes (CD9^+^ EVs) and exosomes (CD63^+^ EVs) released by cells as plating density increased. The O.D. values from the slot blot in **a** are represented as absolute values in **b** and normalized on the cell number in **c**. Notice that the absolute intensities of both CD9 and CD63 tend to increase with increasing cell density (**b**) while the signals decrease when normalized to the number of cells (**c**). This suggests that the number of CD9^+^ and CD63^+^ EVs per cell decreases upon increased cell density. **d**, SUM159 cells expressing CD9-Emerald were imaged by confocal microscopy. Three dimensional volumes were acquired by collecting 22 planes spaced by 500 nm steps in the axial dimension. A representative single plane and the relative YZ profile show that CD9-Emerald is almost exclusively distributed at the cell surface. The white dashed line indicates the plane at which the YZ profile was computed. **e-g**, The concentration of EVs in the supernatants cleared from debris (5,000 x g, 10 minutes) of untransfected (NT) SUM159 and SUM159 expressing CD9-Emerald was measured by Nanoparticle Tracking Analysis (NTA). **e**, The graph (mean ± SEM from three independent experiments) shows the absolute EV concentration (blue line) and the number of EVs released per cell (red line) in the supernatants of SUM159 cells expressing CD9-Emerald. These results follow a similar trend to that of the control (Fig. 1a). Additionally, CD9-Emerald expression neither affects EV concentration (**f**) nor dimension (**g**), and the results reflect the sizes observed for the controls (Fig. 1b-c). Both **f** and **g** show the mean ± SD of three independent experiments. Data were compared using a two-tailed t-test (ns = not significant).

**Figure S2.**
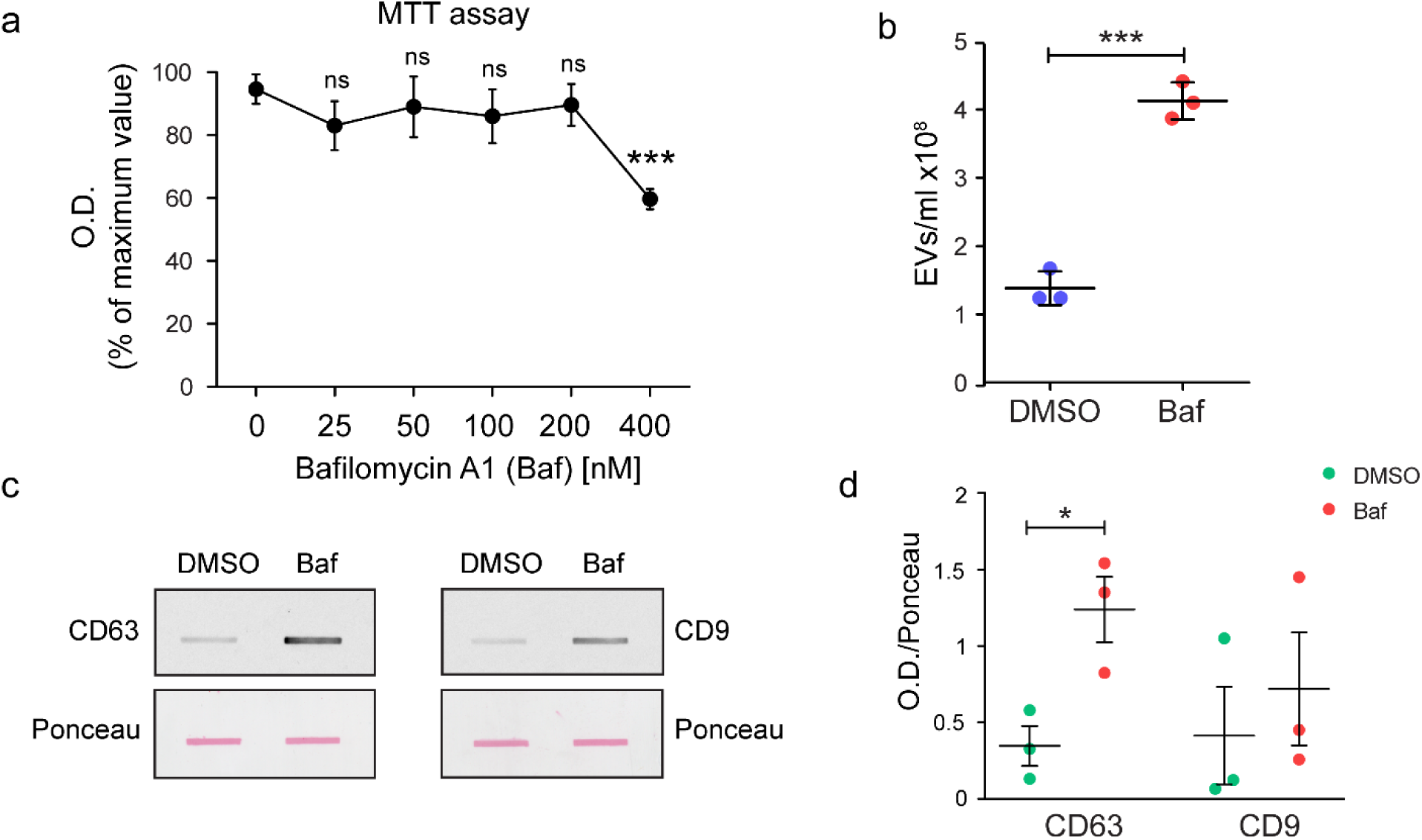
Treatment with Bafilomycin A1 increases the number of EVs retrieved from the supernatant. **a**, Cell viability assay (MTT) was used to define dosages of Bafilomycin A1 (Baf) diluted in Opti-MEM that do not affect the viability of HeLa cells. HeLa cells were treated with either increasing concentrations of the drug (25-400 nM) or with the vehicle DMSO (0 nM) for 24 hours. The graph represents the O.D. values of the MTT metabolite formazan. Bafilomycin A1 did not affect HeLa cell survival at concentrations < 400 nM. Data were compared using a two-tailed t-test (ns = not significant; ^**^ p-value of < 0.0001. 0 nM vs. 25 nM: p = 0.0816; 0 nM vs. 50 nM: p = 0.4081; 0 nM vs. 100 nM: p = 0.1985; 0 nM vs. 200 nM: p = 0.3508). **b**, HeLa cells were cultured in Opti-MEM supplemented with either 50 nM Baf or its solvent, DMSO, for 24 hours. After centrifuging at 5,000 x g for 10 minutes to discard large debris, the cell supernatants were analyzed by Nanoparticle Tracking Analysis (NTA) to quantify the number of particles. The concentration of EVs retrieved from cells treated with Baf (1.4 ± 0.25 × 10^8^; mean ± SD) was significantly increased in comparison to the control (4 ± 0.27 × 10^8^; mean ± SD) across three independent experiments. Slot blots (**c**) and their quantifications (**d**) show that CD63 signal (4.44 ± 1.15; mean ± SEM) increases significantly upon exposure to 50 nM Baf in comparison to CD9 signal (2.99 ± 0.81; mean ± SEM). The respective Ponceau images were used to normalize the slot blot signals. Data were compared using a two-tailed t test where ^*^; p-value < 0.05, ^**^ p-value < 0.001, and ^**^ p-value < 0.0001.

**Figure S3.**
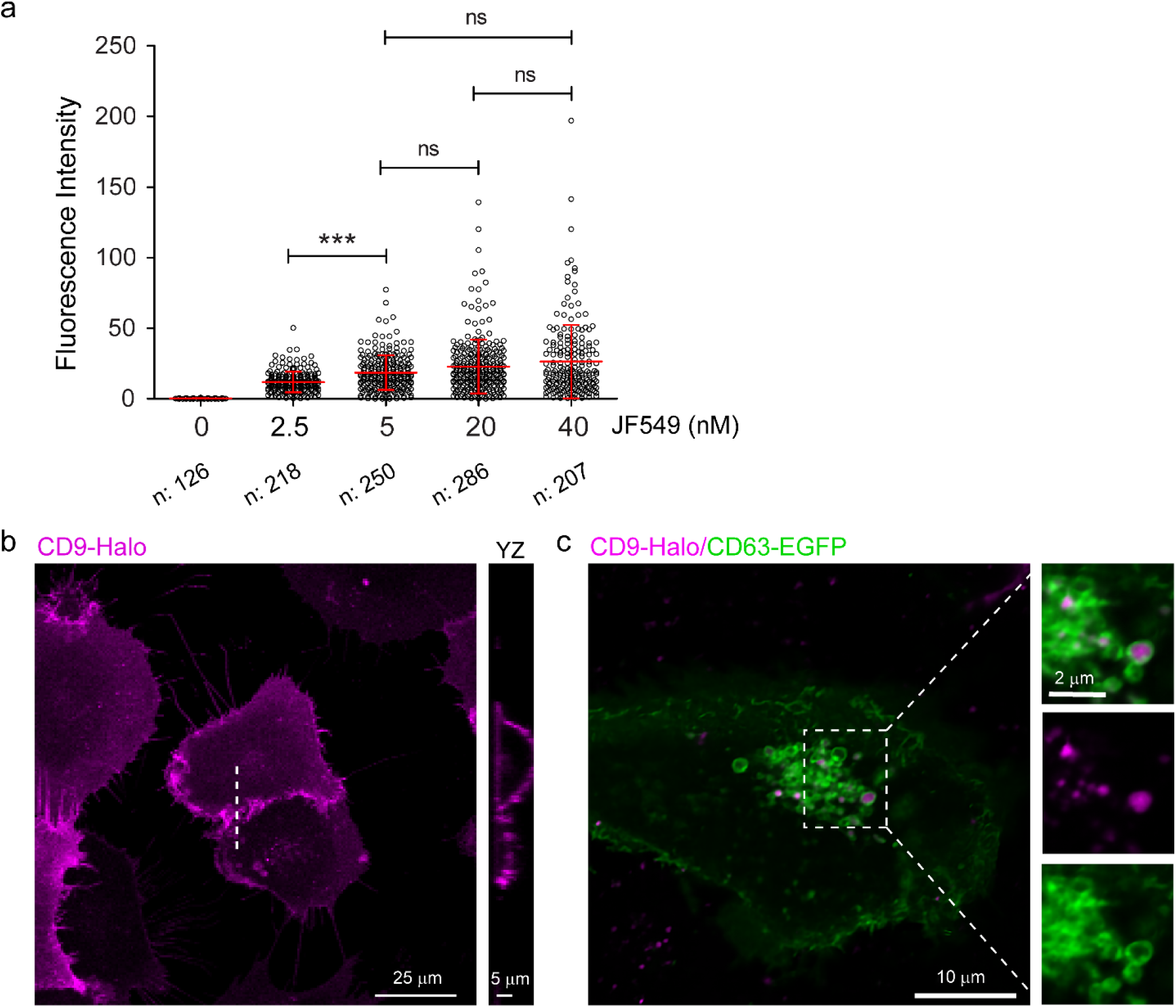
CD9-Halo labelling and distribution. **a**, SUM159 cells stably expressing CD9-Halo were seeded on glass coverslips. Once the cells were adherent and had returned to their typical morphology, they were stained with increasing concentrations of JF549 fluorescent ligand (concentrations indicated in the plot) for 1 hour. Next, the cells were washed in complete media for 2 hours to remove the unbound ligand and subsequently imaged by confocal microscopy. Hoechst staining was used to label the nuclei. The single-cell fluorescent intensity values were measured using the Cell Profiler software package. Incubating the cells with 5 nM JF549 ligand provided a fluorescence intensity that was not significantly different than the ones generated by higher ligand concentrations, suggesting that treatment with 5 nM is sufficient to saturate CD9-Halo. The number of cells analyzed per condition are reported as “n” below the x axis. Data were compared using One-way ANOVA and Tukey’s *post-hoc* test (ns = not significant, ^**^ p-value < 0.0001). **b**, HeLa CD9-Halo cells labelled with 5 nM JF549-Halo as previously described were imaged by confocal microscopy (22 consecutive planes over the Z axis spaced 0.5 μm apart were acquired). A representative image shows that CD9-Halo is mainly distributed over the cell surface as was previously demonstrated for CD9-Emerald chimera (see Fig S1d). The white dotted line indicates the plane at which 3D reconstruction occurred (YZ panel). **c**, HeLa cells expressing CD9-Halo or CD63-EGFP were co-cultured at high density (80% final confluence) for 24 hours. Next, the cells were treated with 50 nM Bafilomycin A1 for an additional 24 hours, stained with 5 nM JF549 for 1 hour as previously described, and imaged by confocal microscopy to assess the distribution of CD9^+^ EVs in the recipient cells. The majority of CD9^+^ ectosomes are enclosed by CD63-labelled organelles, suggesting that they are engulfed in late endosomes and lysosomes.

**Figure S4.**
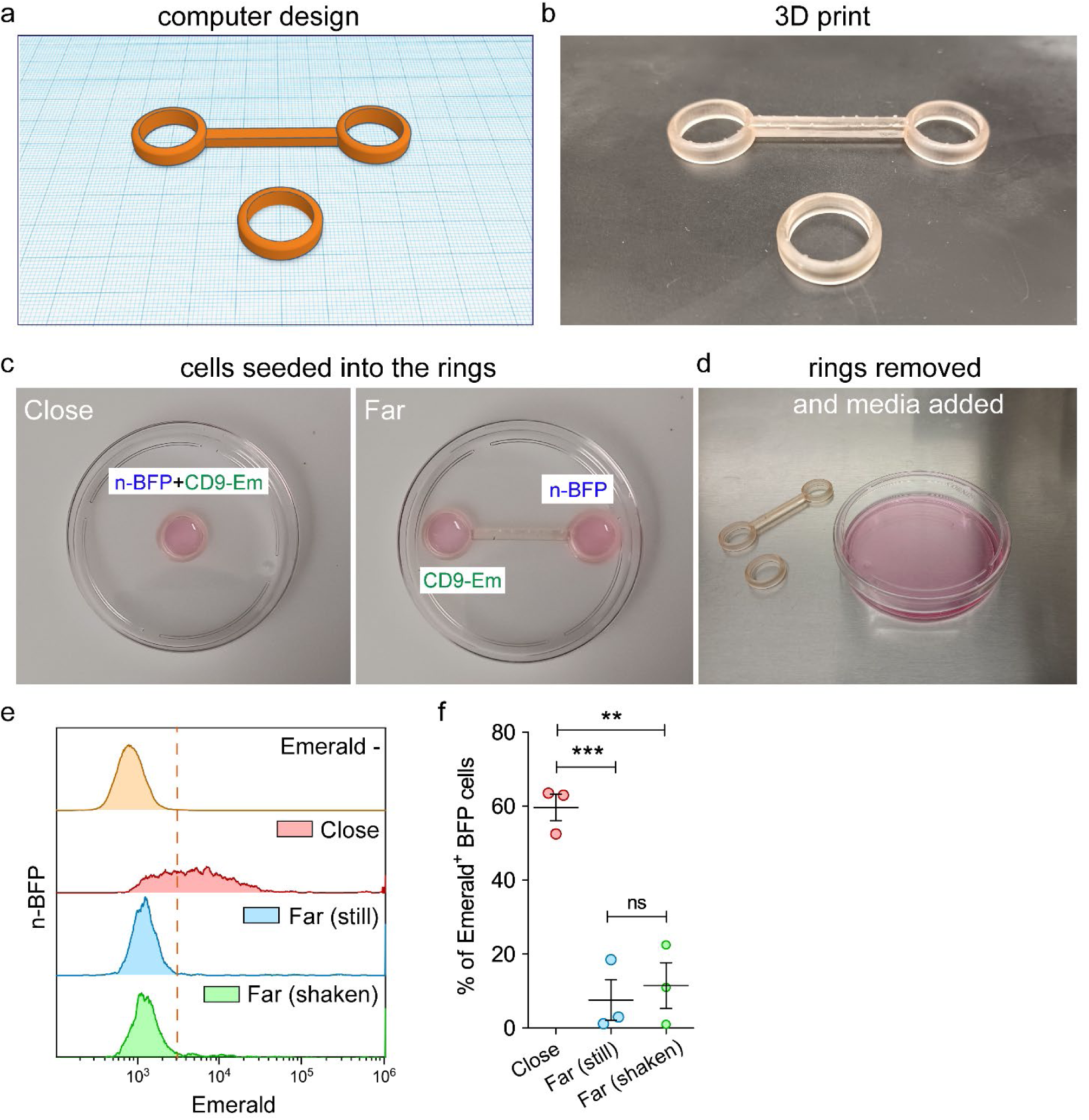
A compartmentalized co-culture system to study the effect of intercellular distance on EV exchange. **a-d**, To define the effect of intercellular distance on the efficiency of EV exchange, we custom designed two co-culture configurations using Tinkercad (Autodesk) software (**a**) and 3D printed those in a biocompatible material (**b**). The “*Close* configuration” was accomplished by placing the single ring in the middle of a 10 cm petri dish. Both donor and recipient cells were plated together at a 1:1 ratio in this compartment (**c**, Close). In comparison, the “*Far* configuration” of cell plating was accomplished using two rings connected by a stem of 4 cm in a 10 cm petri dish. Donor and recipient cells were plated in separate rings (**c**, Far) and the stem ensured consistent distance between cells across experimental replicates. 24 hours post-seeding, the rings were removed, and the cells were cultured for an additional 24 hours in media supplemented with 50 nM Bafilomycin A1 (**d**). **e**, To test the contribution of paracellular fluid flow in EV transport, we set up three co-cultures: one was plated in the *Close* configuration as a control of EV exchange while other two were seeded as *Far* configurations. Of the two *Far* configurations, one co-culture was placed on an orbital shaker at 37°C to promote media movement and simulate the flow of extracellular fluid (*Far* (shaken)) while the other remained stationary (*Far* (still)). After 24 hours, the cells from each condition were detached and analyzed by flow cytometry to measure the percentage of n-BFP^+^/CD9-Emerald^+^ cells. n-BFP^+^ cells seeded without donor cells were used as negative controls to define the minimum value limit for Emerald^+^ signal (yellow distribution and red dashed line). The red distribution accounts for the CD9-Emerald intensities recorded in the n-BFP^+^ cell population cultured in the *Close* configuration, while the blue and the green distributions account for the cells seeded in the *Far* (still) and *Far* (shaken) configurations respectively. The results from three independent experiments are reported in **f**. Data were compared using One-way ANOVA and Tukey’s *post-hoc* test (ns = not significant;^**^ p-value < 0.001 and ^**^ p-value < 0.0001).

**Movie 1**, A confluent co-culture of SUM159 cells expressing CD9-Halo (magenta) and AP2-EGFP (green) was imaged by Lattice Light Sheet Microscopy (LLSM). CD9-Halo was stained by incubating the cells for 1 hour with 5 nM JF549 before imaging. The movie was acquired in sample scan mode and were composed of sequential volumes that corresponded to 40 slices spaced every 350 nm after de-skewing (total volume of 73 × 53 x13 μm). Each slice was exposed to each laser line for 35 ms, resulting in a final frequency of acquisition of 0.3 Hz. The de-skewed stacks were deconvolved using the Lucy-Richardson algorithm (MATLAB, MathWorks) and rendered with a multi-dimension viewer developed in MATLAB environment. The movie is composed of 200 sequential frames mounted at 7 fps. Scale bar is 5 μm. A slice of the movie is shown in Figure 3a. **Dashed inset:** this shows a magnified area of Movie 1 where a single EV is transferred from the CD9-Halo donor to the AP2-EGFP recipient cell. The transfer is completed within the first 24 seconds of the movie. The movie inset is generated at 1 fps and looped, to improve the visualization of the EV exchange. Scale bar is 2 μm. The frames extrapolated from the inset are displayed in Figure 3b. **Solid inset:** this magnified area of Movie 1 shows CD9-Halo EVs stalling at the cell surface of recipient AP2-EGFP cells without been internalized. Puncta of AP2-EGFP, corresponding to clathrin-coated vesicles (CCVs), repetitively surround the EVs for more than 5 minutes without completing an effective internalization. Scale bar is 1 μm. Key frames of this inset are displayed in Figure 3c.

**Movie 2**, A confluent co-culture of SUM159 cells expressing CD9-Halo (magenta) or AP2-EGFP (green) imaged by Lattice Light Sheet Microscopy (LLSM). CD9-Halo was stained by incubating the cells for 1 hour with 5 nM JF549 before imaging. The movie was acquired in sample scan mode and was composed of sequential volumes that corresponded to 40 slices spaced every 350 nm after de-skewing (total volume of 73 × 53 x13 μm). Each slice was exposed to each laser line for 35 ms, resulting in a final frequency of acquisition of 0.3 Hz. The de-skewed stacks were deconvolved using the Lucy-Richardson algorithm (MATLAB, MathWorks) and rendered with a multi-dimension viewer developed in MATLAB environment. The movie is composed of 200 sequential frames mounted at 7 fps. Scale bar is 5 μm. A slice of the movie is shown in Figure 3d. **Dashed inset**: this shows a magnified area of Movie 2 where a single EV is transferred from the CD9-Halo donor to the AP2-EGFP recipient cell. The movie is composed of 200 sequential frames. Scale bar is 2 μm. Key frames of the movie are displayed in Figure 3e.

**Movie 3**, A confluent co-culture of SUM159 cells expressing CD9-Halo (magenta) or AP2-EGFP (green) imaged by Lattice Light Sheet Microscopy (LLSM). CD9-Halo was stained by incubating the cells for 1 hour with 5 nM JF549 before imaging. The movies were acquired in sample scan mode and were composed of sequential volumes that corresponded to 40 slices spaced every 350 nm after de-skewing (total volume of 73 × 53 x13 μm). Each slice was exposed to each laser line for 35 ms, resulting in a final frequency of acquisition of 0.3 Hz. The de-skewed stacks were deconvolved using the Lucy-Richardson algorithm (MATLAB, MathWorks) and rendered with a multi-dimension viewer developed in MATLAB environment. The movies shown are composed of 133 and 177 sequential frames respectively mounted at 7 fps. Scale bar is 4 μm. The white arrows indicate filopodia of CD9-Halo cells that fragmented in the extracellular space into EVs.

## Notes

### Competing Interest Statement

The authors have declared no competing interest.

